# METANet: A supervised ensemble learning framework for reconstructing direct and functional tissue-specific transcription factor networks

**DOI:** 10.1101/2025.10.10.681634

**Authors:** Wooseok J. Jung, Sandeep Acharya, Daniel P. Ruskin, Shu Liao, Vaha Akbary Moghaddam, Zolboo Erdenebaatar, Michael R. Brent

## Abstract

**Motivation:** Reconstructing tissue-specific transcription factor (TF) networks remains challenging. TF motif-based methods often lack functional validation, while expression-based methods struggle to distinguish direct binding from indirect regulation. Integration of diverse data types is necessary to accurately prioritize functional targets directly bound by TFs across human tissues.

**Results:** We introduce METANet, a supervised ensemble learning framework that combines TF motifs, cis-regulatory element activity, and linear and non-linear expression-derived features to predict TF binding. Applied to 36 human tissues, METANet significantly outperforms established methods in identifying direct, functional targets of TFs validated by ChIP-seq and gene ontology. Furthermore, METANet captures tissue-specific regulation comparable to existing methods, allowing the identification of reproducible gene-trait associations.

**Availability and Implementation:** All code and network maps are freely available at Zenodo https://doi.org/10.5281/zenodo.17309371.

## 1 INTRODUCTION

A transcription factor (TF) network map is a directed graph in which nodes represent TFs and genes, and edges link TFs to the genes they directly regulate. Numerous algorithms have been developed for mapping TF networks from a variety of data types, and these methods generally fall into two broad categories. The first category uses TF binding evidence, such as DNA sequence motifs (Janky, et al., 2014) or TF ChIP-seq (Gerstein, et al., 2012), often combined with promoter and enhancer annotations to define regulatory elements. Motif data describe locations where a TF could bind and TF ChIP-seq data reveal the genomic regions where a TF actually binds. However, binding evidence alone cannot confirm whether a TF functionally affects the transcription rates of its target genes. For instance, many TF-bound motifs have no apparent effect on gene transcription (Cusanovich, et al., 2014; Slattery, et al., 2014; White, et al., 2013). Furthermore, most motifs for a TF are not bound by the TF (Dror, et al., 2015; Wang, et al., 2012), and many sites where a TF binds lack motifs (Horton, et al., 2023; Worsley Hunt and Wasserman, 2014).

Another class of methods uses gene expression measurements under baseline conditions or after perturbations of TF expression (Abid and Brent, 2023; Haynes, et al., 2013; Huynh-Thu, et al., 2010; Kang, et al., 2018; Langfelder and Horvath, 2008; Skok Gibbs, et al., 2022). While perturbation-response data can reveal the functional effects of TFs, they do not confirm direct binding. Moreover, expression-based methods are prone to capturing indirect effects, as correlations between TF and target gene (TG) expression may arise from transitive effects such as shared upstream regulators or mediated interactions (Feizi, et al., 2013; Mercatelli, et al., 2020; Xiao, et al., 2022). To address these limitations, prior studies have established that integrating sequence-specific motifs as priors with gene expression data significantly improves regulatory network inference compared to expression-based inference alone (Greenfield, et al., 2013; Roy, et al., 2013; Siahpirani and Roy, 2017).

While many regulatory relationships are shared across tissues, some TF-TG interactions are context-specific. TF network maps inferred from gene expression and binding data have been shown to vary significantly across cell types and tissues (Gamazon, et al., 2018; Haigis, et al., 2019; Melé, et al., 2015; Sonawane, et al., 2017). Our group has previously mapped TF networks in *Saccharomyces cerevisiae* and *Drosophila melanogaster* using both TF binding and gene expression data, successfully capturing direct and functional targets of TFs (Abid and Brent, 2023; Kang, et al., 2018). However, these mapping efforts focused on a single, aggregate context and did not address regulatory variation across tissues, which limits their utility for understanding context-specific gene regulation in human tissues. Most widely used approaches for tissue-specific TF network mapping remain limited in their ability to capture direct and functional targets. For example, TF motif-based approaches using cis-regulatory element (CRE) activity levels across tissues can suggest tissue-specific binding potential, but do not confirm functional relationships (Marbach, et al., 2016), while correlation-based networks fail to account for the physical binding of the TF in the target gene’s CREs (Pierson, et al., 2015; Saha, et al., 2017). One notable exception is (Sonawane, et al., 2017) which uses PANDA, combining TF motifs and protein-protein interactions (PPI) with tissue-specific expression data in a message-passing framework, to map 38 tissue-specific TF networks. Building on our prior work, we introduce the METANet framework, a supervised machine learning approach to predict tissue-specific direct and functional TF-TG regulatory relationships. Unlike PANDA’s unsupervised approach, METANet uses supervised learning which allows the model to learn directly from TF ChIP-seq-based binding signals. Using tissue-specific gene expression data and TF network maps derived from (Marbach, et al., 2016) as predictor variables, METANet predicts the probability that a TF binds a target gene in TF-ChIP-seq data the model has not seen.

This paper makes four contributions. We provide a comprehensive resource of human tissue-specific TF network maps across 36 tissues, encompassing ∼237 TFs and ∼12,150 protein-coding genes. We evaluate the ability of our TF network maps to capture direct and functional targets of TFs using four evaluation metrics and compare our performance to the networks proposed by Sonawane et al. We also compare to the original networks from Marbach et al (which does not use gene expression data). Across 36 tissues, we show that METANet better captures both the direct and functional targets of TFs compared to benchmark network maps. We also found that the tissue-specificity of METANet is comparable to that of the network maps from Marbach et al and Sonawane et al. To explore potential applications of METANet, we input it to FISHNET (Acharya, et al., 2025), a network-driven gene prioritization method, using transcriptome-wide association (TWAS) summary statistics for traits associated with cardiovascular risk from the Long Life Family Study (LLFS) cohort. METANet identified replicated gene-trait associations that were missed when other networks were passed as inputs to FISHNET. Furthermore, METANet identified regulatory mechanisms that may mediate the effect of known trait-associated variants.

## 2 MATERIALS AND METHODS

### 2.1 METANet

METANet combines five distinct sets of regulatory edge features using XGBoost: a tissue-specific motif-based feature (Marbach) and four expression-based features derived from LASSO and BART regression on both tissue-specific and tissue-aggregate contexts (Fig. 1A). The Marbach scores are based on motif occurrences weighted by tissue-specific cis-regulatory element activity. LASSO and BART scores are derived from regressing each gene’s mRNA levels on those of all TFs using either tissue-specific samples or samples aggregated across all tissues. These five weighted networks form a set of five features that are trained with binary labels extracted from binding experiments: 1 when there is evidence of direct binding within the cis-regulatory elements of a gene and 0 otherwise. We trained the model using 10-fold nested cross-validation (CV), stratified based on the distribution of binding labels. A new XGBoost model was trained for each outer fold, using the remaining 9 folds as training data and the remaining fold as test data. We further divided each training fold using 5-fold CV for hyperparameter tuning; the optimal parameters were used for training an XGBoost model with all the training data to generate predictions for the hold-out test fold. A TF was not allowed to regulate itself.

**Fig. 1.**
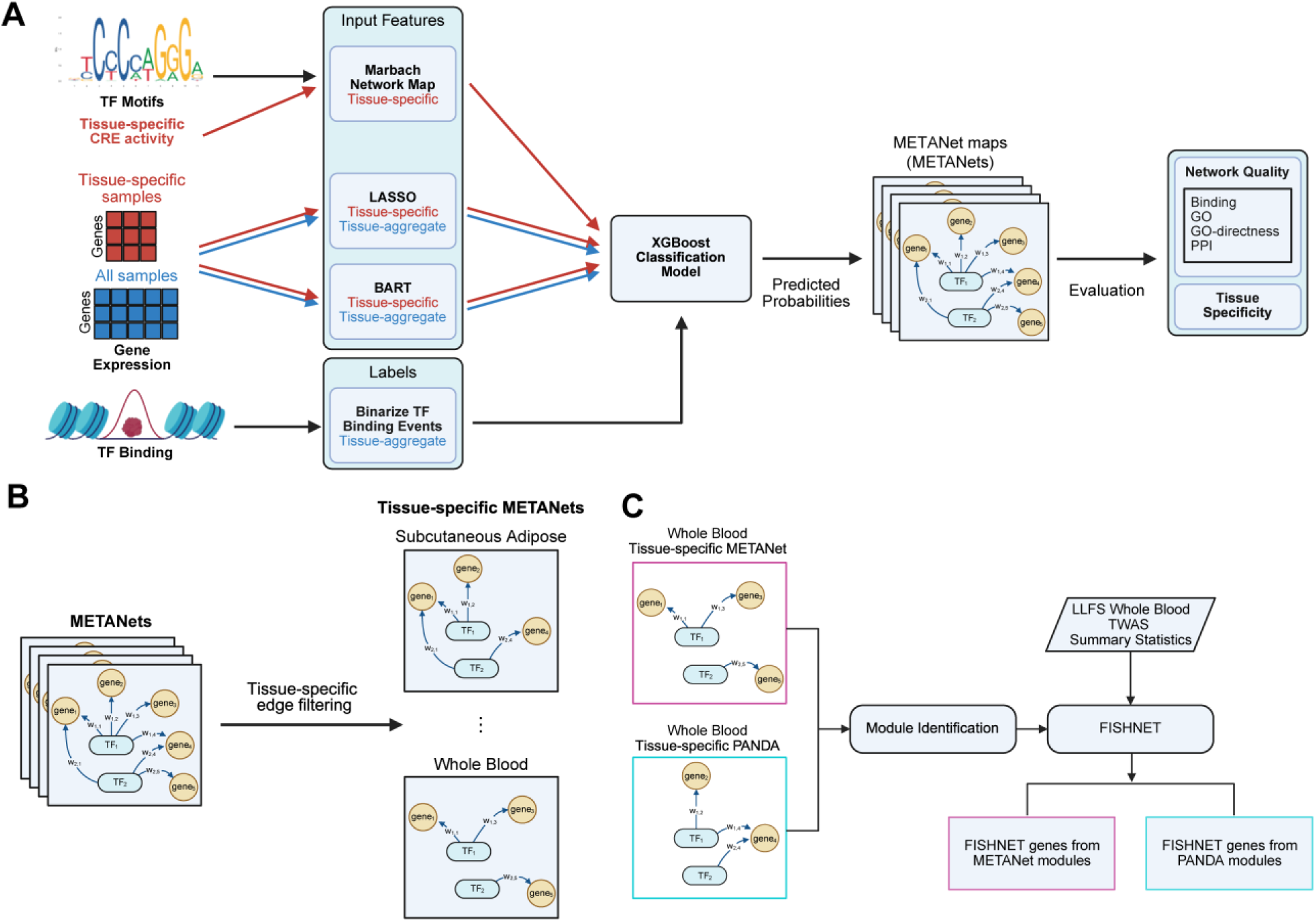
METANet framework overview. (**A**) For each tissue, METANet combines evidence scores from tissue-specific TF network maps from Marbach et al. 2016, tissue-specific and tissue-aggregate LASSO and BART regression models from GTEx expression data, and binarized TF binding labels using XGBoost. The predicted probabilities from the XGBoost model are used as edge scores in our tissue TF network maps. Red arrows indicate tissue-specific data while blue arrows indicate tissue-aggregate data. The predicted METANet maps (METANets) are evaluated for network quality and tissue specificity. (**B**) Tissue-specific METANets are inferred from the predicted METANets by filtering for tissue-specific edges. (**C**) Modules identified from whole blood-specific METANet and PANDA networks are used in FISHNET with TWAS summary statistics from the LLFS to identify FISHNET genes.

### 2.2 Construction of Binding Labels

For each TF, we identified high-confidence (FDR q-value ≤ 0.01) ChIP-seq binding peaks from REMAP 2020 (Cheneby, et al., 2020) within FANTOM5 promoter and enhancer regions (Andersson, et al., 2014; Consortium, et al., 2014). Gene-level scores were calculated by summing the confidence scores of non-overlapping peaks across all CREs associated with a target gene. We constructed binary labels representing TF-gene binding events by assigning the top 10% of highest-scoring genes as positive labels. All remaining TF-gene pairs were labeled as negative.

### 2.3 Construction of Expression-based Features

We downloaded bulk RNA-seq read counts and transcripts per million (TPM) data from the Genotype-Tissue Expression (GTEx) consortium V8 release (Aguet, et al., 2020). The GTEx read count data for protein-coding genes was filtered to remove lowly expressed genes. Specifically, genes with ≤ 3 counts per million in ≥ 98.5% of samples were removed.

We constructed the gene expression-based features by training two regression algorithms, LASSO and Bayesian Additive Regression Trees (BART), to predict the expression level of each gene from the expression levels of all TFs (Abid and Brent, 2023; Chipman, et al., 2010; Haynes, et al., 2013; Kang, et al., 2018). LASSO (Least Absolute Shrinkage and Selection Operator) is a linear regression method with L1 regularization. BART is a non-parametric sum-of-trees model that captures complex, non-linear relationships through an ensemble of regression trees. Features for a given TF-gene pair reflect the effect of the TF’s expression level on the gene’s predicted expression level in the trained models. For example, the feature extracted from a LASSO model is the learned coefficient of the TF’s expression level. Both algorithms are run on the filtered tissue-specific gene expression TPM data and, separately, on the aggregated TPM data from all GTEx tissue samples, yielding 4 regression features for each TF-gene pair.

### 2.4 Marbach Networks

We downloaded 394 cell-type- and tissue-specific networks from Marbach et al. (Marbach, et al., 2016). We formed 36 tissue networks by merging closely related networks by taking the graph union (union of the node and edge sets while retaining the maximum weight for each edge). To guide this merging process, we adopted the tissue-to-cell mapping established by Marbach et al. as the baseline reference for 13 GTEx tissues. We then extended this by manually curating mappings for the remaining 23 tissues using annotations from the BRENDA tissue ontology (Gremse, et al., 2011) (see Supplementary Table S3).

### 2.5 PANDA Networks

For benchmarking, we obtained PANDA networks for the same 26 tissues from Sonawane et al. (Sonawane, 2017; Sonawane, et al., 2017). We processed both fully connected and tissue-specific PANDA networks identically to METANets, restricting analyses to protein-coding genes and the common set of transcription factors to ensure fair comparison (see Supplementary Methods).

### 2.6 Network Quality Evaluation

#### 2.6.1 Binding metric

We calculated the percentage of top-scoring TF-TG edges supported by TF binding data at different thresholds. Top-scoring edges were defined by sorting edges by the edge score in descending order. Thresholds were scaled to the number of target genes per TF. This is an average, so different TFs have different numbers of target genes at each threshold. A higher percentage of binding support reflects greater enrichment of direct edges in the network map. The random expectation is the probability that a randomly selected TF-TG edge is supported by the binding data.

#### 2.6.2 GO metric

We performed GO enrichment analysis for each TF’s targets using GO-Term-Finder v0.86 (Boyle, et al., 2004). To focus on terms associated with specific biological processes, rather than extremely generic terms like ‘biosynthetic process’, we excluded terms with number of annotated genes below 5 or exceeding 300. Each TF’s score was the maximum –log P-value over all GO terms. If a TF’s target set showed no significant enrichment for any GO biological process, we assigned the TF a P-value of 1 (equivalent to –log P = 0). The final network-level score was taken as the median of these scores across all TFs. We did these analyses for each threshold, as in the binding metric, but without excluding TFs that did not have binding data available.

#### 2.6.3 GO-directness metric

To determine whether improvements on the GO evaluation metric came at the cost of including indirect edges, we calculated, for each TF, the percentages of binding support for edges that matched the GO term with the highest minus log P-value. For each threshold, we reported the average percentage of binding support across TFs. The random expectation is identical to that of the Binding metric.

#### 2.6.4 PPI metric

First, we calculated the Jaccard similarity between the sets of target genes of each (TF, TF) pair; the Jaccard similarity is 1 when the target sets are the same and 0 when none of the targets in the sets are shared. Next, we sorted (TF, TF) pairs by Jaccard similarity and calculated the percentage of the top 100 pairs that are supported by physical interactions from the protein-protein interaction (PPI) STRING database (Szklarczyk, et al., 2023). The database has 477,903 interactions predicted with high confidence (SRTING score ≥ 0.7). A higher percentage of PPI support means that (TF, TF) pairs are likely to work as a part of a known physical protein complex. We did these analyses for the TF-TG edges at thresholds of 50 and 100 targets per TF. The random expectation is the probability that a randomly selected pair of TFs present in a tissue network is supported by the PPI database. This probability was computed separately for each tissue and then averaged across the 36 tissues.

#### 2.6.5 Empirical Null

To establish an empirical null to evaluate network quality, we generated 50 permuted networks by randomly shuffling the fully connected METANet edge scores. Each permuted network was subjected to the same cutoff thresholds as the real networks and evaluated across all four metrics, providing a consistent null distribution.

### 2.7 Tissue Specificity Evaluation

#### 2.7.1 Tissue-Specific Edge Filtering

Tissue-specific regulatory edges were identified using the interquartile range (IQR)-based filtering specificity score described by Sonawane et al. (Sonawane, et al., 2017). Edges with a normalized specificity score > 2 were retained as tissue-specific (see Supplementary Methods).

#### 2.7.2 Validation of Tissue Specificity using eQTL Data

We validated tissue specificity by intersecting network edges with GTEx V8 cis-eQTL in target gene regulatory regions (The GTEx Consortium, 2020). We calculated ‘eQTL support’ and ‘eQTL count’ metrics (see Supplementary Methods). Significance was assessed using a permutation test (10000 iterations) on the sum of ranks for matching tissue pairs, and networks were compared using the Wilcoxon signed-rank test.

### 2.8 Gene-trait Association Analysis using FISHNET

We evaluated the biological relevance of METANets using FISHNET, a network-based gene prioritization tool (Acharya, et al., 2025). FISHNET is designed to prioritize genes that exhibit suggestive association signals but are more likely than expected by chance to replicate across independent datasets. It operates on the intuition that genes with low p-values purely by chance are distributed randomly, whereas true associations tend to cluster within densely connected network modules and share common biological functions. Using whole blood tissue-specific METANets and PANDA networks as inputs, we partitioned networks into modules using the M1 modularity optimization algorithm (Tomasoni, et al., 2020). We integrated TWAS p-values for 11 cardiovascular traits from the Long Life Family Study (LLFS) and assessed replication using summary statistics from the Framingham Heart Study (FHS).

## 3 RESULTS

### 3.1 Overview of TF Network Map Reconstruction

To map tissue-specific transcription factor (TF) regulatory networks, we developed the METANet framework, a supervised learning approach that integrates TF motif data, gene expression-derived features, and TF binding evidence. For each of 36 human tissues, we trained an XGBoost model (Chen and Guestrin, 2016) to predict whether a TF binds within the cis-regulatory elements (CREs) of a gene, using ChIP-seq data from REMAP 2020 (Cheneby, et al., 2020) as ground truth (Fig. 1). Each instance in the model corresponds to a candidate TF-target gene (TG) pair and was represented by five biologically motivated features: (1) a motif-based regulatory score from the tissue-specific Marbach network (Marbach, et al., 2016), which combines TF motif occurrences with tissue-specific cis-regulatory element (CRE) activity; (2–5) four expression-derived features capturing both linear and nonlinear TF-TG relationships. The features were derived by regressing the expression of each gene on the expression of all TFs using both LASSO (Tibshirani, 1996) and Bayesian Additive Regression Trees (BART) (Chipman, et al., 2010). We extracted from the model for each gene measures of the relatedness of each TF’s expression to the gene’s expression (see Methods for details). Both regression algorithms were applied to both tissue-specific and tissue-aggregated RNA-seq data from GTEx (The GTEx Consortium, 2020) (Fig. 1). We used 10-fold nested cross-validation to train and evaluate each model. Predicted probabilities from test folds were used as edge scores in the resulting METANet maps (METANets), linking TFs to their likely direct functional targets. In total, we mapped the TF networks of 36 human tissues, encompassing an average of 237 TFs and 12150 genes.

### 3.2 METANets Capture Direct and Functional Targets of TFs

We compared METANets against the motif-based Marbach networks, constructed using motif occurrences and tissue-specific CRE activity levels (Marbach, et al., 2016), and PANDA networks from Sonawane et al., built using motifs and expression via an unsupervised approach (Sonawane, et al., 2017), across 26 common tissues. For the PANDA networks, we used the fully connected PANDA output networks from Sonawane et al. prior to filtering for tissue-specific edges. All networks we evaluated include continuous edge scores, from which discrete networks can be created by thresholding and retaining only edges above the chosen cutoff. To enable fair comparisons across tissues, we applied thresholds scaled to the average number of targets per TF (e.g. top 20000 edges for 200 TFs implies 100 targets per TF), and we examined how evaluation results changed as more, lower-scoring edges were included.

To assess network quality, we used four complementary metrics: Binding, Gene Ontology (GO), GO-directness, and protein-protein interaction (PPI). To provide an empirical null expectation for each metric, we generated 50 random networks by permuting the edge scores of METANets. These random networks were also subjected to the same cutoff thresholds as other networks before evaluation.

### Binding

The Binding metric is the fraction of edges supported by binding data showing that the TF binds in one or more of the target’s CREs, sourced from the FANTOM5 database (Andersson, et al., 2014; Consortium, et al., 2014). METANets significantly exceeded both Marbach and PANDA (paired *t*-tests, P<0.001 for each; Fig. 2A) and far exceeded the random expectation (*t*-test, P=1.8 × 10^−32^), indicating stronger enrichment for directly bound targets.

**Fig. 2.**
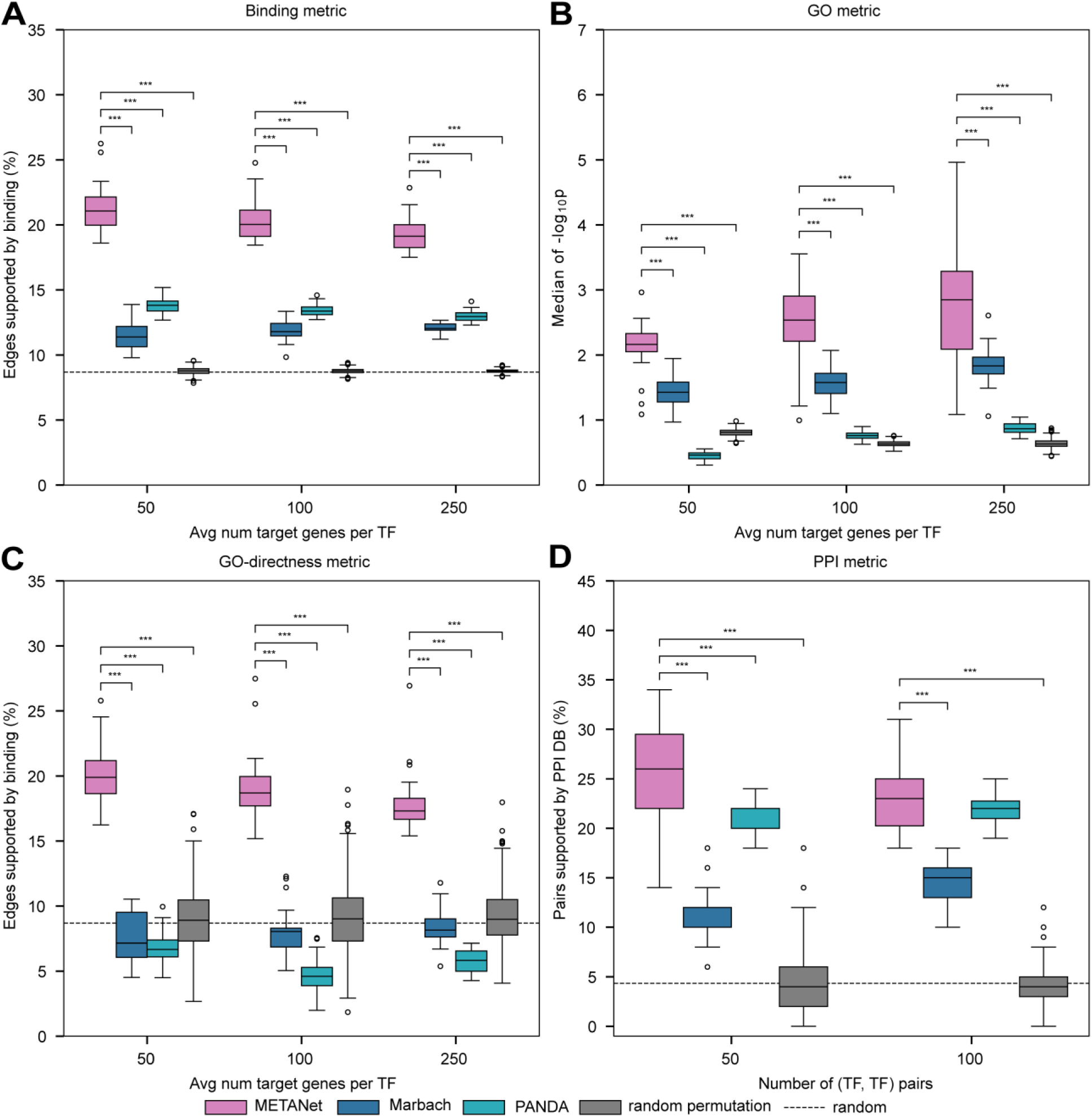
Evaluation of network quality. Performance of METANets, Marbach networks, and PANDA networks on 26 tissues for which all three networks were available. Pink: METANets; Blue: Marbach network maps; Cyan: PANDA network maps; Gray: random expectation based on permutations of METANets; black dashed: random expectation. (**A**) Binding metric (defined in main text) calculated at different edge-score thresholds. METANets outperform all other network maps. Random expectation is the probability that a randomly selected TF-TG edge will be supported by binding data. (**B**) GO metric (defined in main text). METANets outperform all other network maps and the random expectation. Random permutation is the GO metric for 50 METANets in which edge scores were randomly permuted. (**C**) GO-directness metric (defined in main text). METANets significantly outperform all other network maps. Random expectations include the GO-directness evaluations of the GO results of the 50 permuted network maps generated in (B) and the binding metric random expectation. (**D**) PPI metric (defined in main text). METANets significantly outperform all other network maps. The random expectation is the probability that a randomly selected (TF, TF) pair will be supported by PPI data. Significance bars only drawn for significant paired *t-*tests with METANets. \**P≤*0.05, \*\**P≤*0.01, \*\*\**P≤*0.001.

### Gene Ontology (GO) enrichment

To compute the GO metric, we carried out over-representation analysis of GO biological process annotations among the targets of each TF. Each TF’s score was the maximum –log P-value over all GO terms. If a TF’s target set showed no significant enrichment for any GO biological process, we assigned the TF a P-value of 1 (equivalent to –log P = 0). The final network-level score was taken as the median of these scores across all TFs. METANets outperformed Marbach and PANDA (both P<0.001; Fig. 2B) and surpassed the random permutation (P<0.001; see Methods), showing greater functional coherence among the predicted targets of each TF.

### GO-directness

The GO-directness metric is the fraction of targets annotated with each TF’s most significant GO term that are also supported by ChIP-seq binding. This metric penalizes networks that achieve high GO scores by including many, functionally related, but indirect, targets. METANets retained significantly more binding support than both Marbach and PANDA (P<0.001; Fig. 2C) and the random permutations (P<0.001).

### Protein-protein interaction (PPI)

The PPI metric is the fraction of top-ranked (TF, TF) pairs, based on target set similarity, that are supported by high-confidence physical protein-protein interactions in the STRING database (confidence score ≥ 0.7) (Szklarczyk, et al., 2017). METANets outperformed Marbach (P<0.001) and PANDA (P<0.05), recovering biologically corroborated physical TF-TF interactions (Fig. **2**D).

Together, these results show that METANets are enriched for direct and functionally coherent TF-TG interactions relative to other networks.

### 3.3 Combining Motif and Expression Features Improves Network Mapping

We next evaluated the contribution of TF motif- and gene expression-derived features by comparing METANets to the Marbach networks, which are built using only motifs, and to ETANets, which we built in the same way as METANets but not including the edge score from Marbach networks, using only expression data across all 36 tissues. METANets significantly outperformed both Marbach and ETANet in the Binding metric (P<0.001), indicating the value of integrating both information sources (Fig. 3A). Interestingly, ETANet performed best in the GO metric, suggesting that gene expression alone captures broader functional coherence (Fig. 3B). However, METANets significantly outperformed ETANets in the GO-directness metric (P<0.001), indicating that many of ETANets’ additional functionally coherent targets are likely indirect (Fig. 3C). Thus, METANets provide a more accurate map of direct, functional TF-TG interactions than both the Marbach networks and the PANDAs networks.

**Fig. 3.**
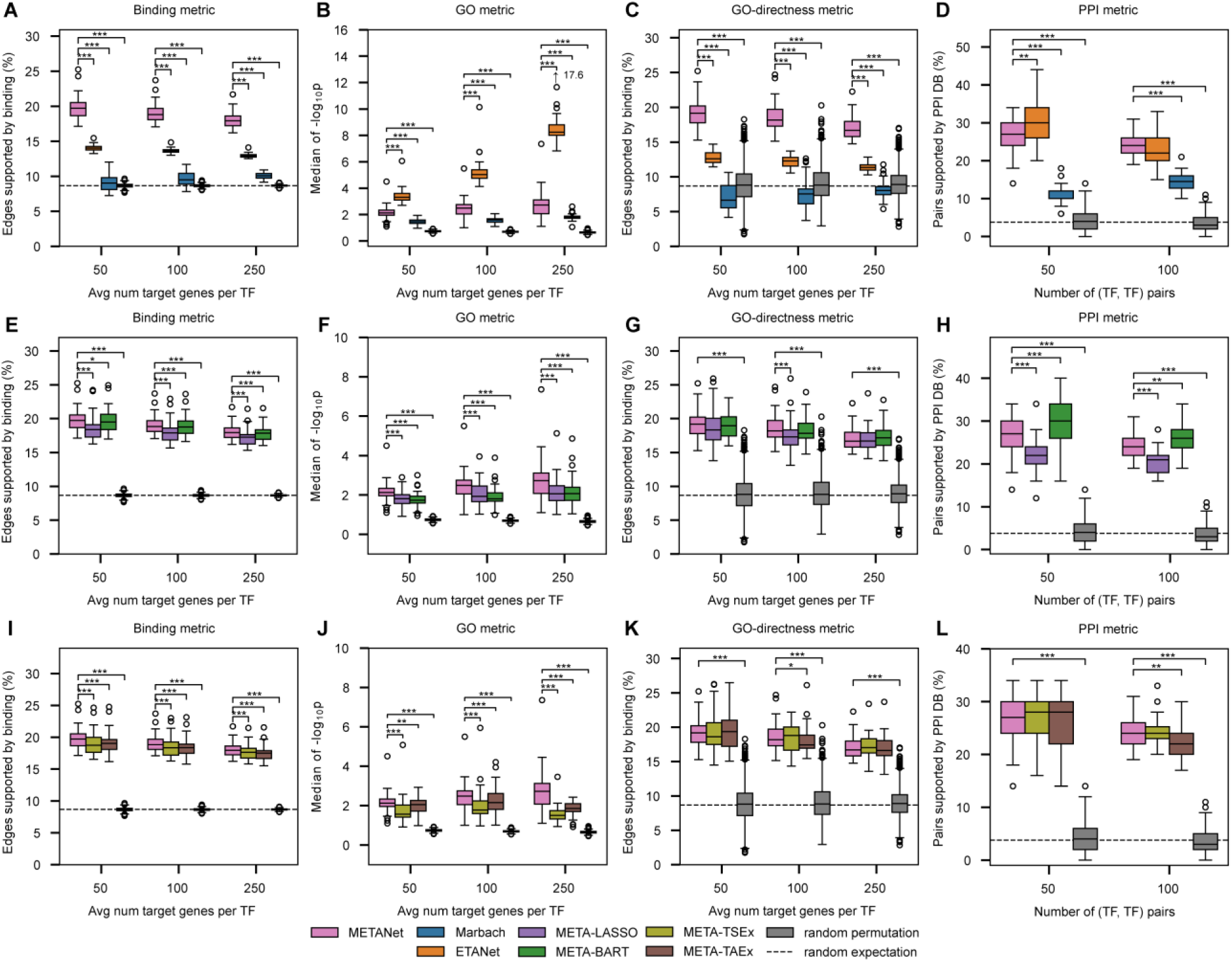
Network quality evaluations against ablated networks. (**A-B**) Performance of ss and the Marbach networks (motif-only) and ETANets (expression-only) on 36 tissues. (**A**) Binding metric. METANets significantly outperformed both ETANets and Marbach networks. (**B**) GO metric. ETANets outperformed METANets and Marbach networks. (**C**) GO-directness metric. METANets outperformed ETANets, predicting more accurate TF network maps of direct and functional TF-TG interactions. (**D**) PPI metric. ETANets outperformed Marbach networks in both thresholds but were comparable to METANets for top 100 (TF, TF) pairs. **(E-H)** Performance of METANets and ablated TF network maps META-LASSO and META-BART. (**E**) Binding metric. METANets and META-BART significantly outperformed META-LASSO, while METANets and META-BART performed comparably. (**F**) GO metric. METANets outperformed both ablated TF network maps. (**G**) GO-directness metric. All three TF network maps significantly outperformed the random expectation based on 50 permutations of METANets. METANets are comparable with both ablated network maps. (**H**) PPI metric. META-BART outperformed both METANets and META-LASSO. **(I-L)** Performance of METANets and ablated TF network maps META-TSEx and META-TAEx. (**I**) Binding metric. METANets significantly outperformed both ablated TF network maps. (**J**) GO metric. METANets outperformed both ablated TF network maps. (**K**) GO directness metric. All three TF network maps significantly outperformed the random permutation and METANets were comparable with both ablated network maps. (**L**) PPI metric. All three network maps performed similarly with and outperformed the random permutation. Significance bars only drawn for significant paired *t-*tests with METANet. \**P≤*0.05, \*\**P≤*0.01, \*\*\**P≤*0.001.

### 3.4 Linear and Nonlinear Expression Features are Synergistic

To assess whether combining linear (LASSO) and nonlinear (BART) expression-derived features improved network quality, we compared METANets against ablated versions using only one feature type: META-LASSO and META-BART. In the Binding metric, METANets and META-BART outperformed META-LASSO. Compared to each other, METANets and META-BART performed comparably (Fig. 3A). In the GO metric, METANets significantly outperformed both ablated network maps (Fig. 3B), suggesting that leveraging both LASSO and BART features better captures functionally coherent relationships. GO-directness was similar across models (Fig. 3D), consistent with gains in functional targets without loss of direct targets.

### 3.5 Combining Tissue-specific and Tissue-aggregate Expression Features is Helpful

To assess the contribution of expression-derived feature specificity, we compared METANets to two ablated variants. META-TSEx retains the motif-based Marbach feature and tissue-specific expression features and excludes tissue-aggregate expression features (LASSO and BART trained on all samples pooled across tissues). Conversely, META-TAEx retains the Marbach feature and tissue-aggregate expression features, but not the tissue-specific expression features. In the Binding and GO metrics, METANets outperformed both ablated models (Fig. 3E, F). In the GO-directness metric, METANets significantly outperformed META-TAEx at one threshold (100 targets per TF) (Fig. 3G). Together, these results show that combining both tissue-specific and tissue-aggregate expression features enables METANets to identify substantially more functionally coherent targets of TFs without sacrificing direct binding support.

### 3.6 METANets are Tissue Specific and Comparable to Existing Networks

We evaluated tissue specificity using tissue-specific expression quantitative trait loci (eQTLs) from GTEx (Consortium, 2020). For each edge in each METANet map, we queried whether at least one tissue-specific eQTL overlapped the TF’s ChIP-seq peaks in the promoter or annotated enhancers of the target gene. We defined two complementary metrics:

1. eQTL support: Fraction of edges supported by at least one tissue-specific eQTL.
2. eQTL count: Sum, across all edges, of the number of tissue-specific eQTLs supporting each edge.

To assess a network’s tissue-specificity, we performed a permutation test. First, we calculated an observed statistic *S*_*obs*_ by ranking all 36 networks within each tissue and summing the ranks of the “matching” pairs (e.g., the rank of the liver network in the liver tissue). A smaller sum indicates better overall specificity. Next, to generate a null distribution, we performed 10000 permutations. In each permutation, we randomly shuffled the network labels before summing the ranks of the newly assigned matching pairs to obtain the permutation statistic *S*_*perm*_. This process simulates the null hypothesis that there is no true correspondence between a network and its native tissue. The empirical p-value was then calculated as the proportion of permutations with *S*_*perm*_ ≤ *S*_*obs*_.

### 3.7 METANets are as tissue specific as the Marbach networks

We compared the tissue specificity of top-scoring edges in METANets and the Marbach networks. METANets showed significant tissue specificity compared to the null in both eQTL support and eQTL count (P<0.05) at higher average targets per TF thresholds (≥100 targets per TF) (Table 1). The Marbach networks showed similar patterns, and the Wilcoxon signed-rank test revealed no significant difference in the distributions of matching tissue ranks between METANets and Marbach networks in tissue specificity (Table 3). Thus, METANets achieve comparable tissue specificity while outperforming the Marbach networks in all network quality metrics.

**Table 1.** Tissue specificity evaluation of METANets and Marbach against the null across 36 tissues. Permutation test p-values (and the sum of observed matching ranks) against the distribution of 10000 permutation sums. METANets showed significant tissue specificity compared to the null in eQTL support at average 100 and 250 targets per TF thresholds. The Marbach networks showed significant tissue specificity in eQTL support at all thresholds. METANets showed significant tissue specificity in eQTL counts at average 50 and 100 targets per TF thresholds. The Marbach networks show significant tissue specificity in eQTL counts at average 250 targets per TF only. See also Supplementary Fig. S1.

| Empirical test for tissue-specificity for 36 tissues |  |  |  |
| --- | --- | --- | --- |
| eQTL support | Average number of targets per TF |  |  |
|  | 50 | 100 | 250 |
| METANets | 0.498 (666) | **0.027 (597)** | **0.022 (613)** |
| Marbach | **0.003 (567)** | **$2 \times 10^{-4}$ (554)** | **$6 \times 10^{-4}$ (537)** |
| eQTL counts | Average number of targets per TF |  |  |
|  | 50 | 100 | 250 |
| METANets | 0.106 (604) | **0.003 (535)** | **0.007 (563)** |
| Marbach | 0.119 (611) | 0.173 (623) | **$3.4 \times 10^{-3}$ (537)** |

**Table 2.**
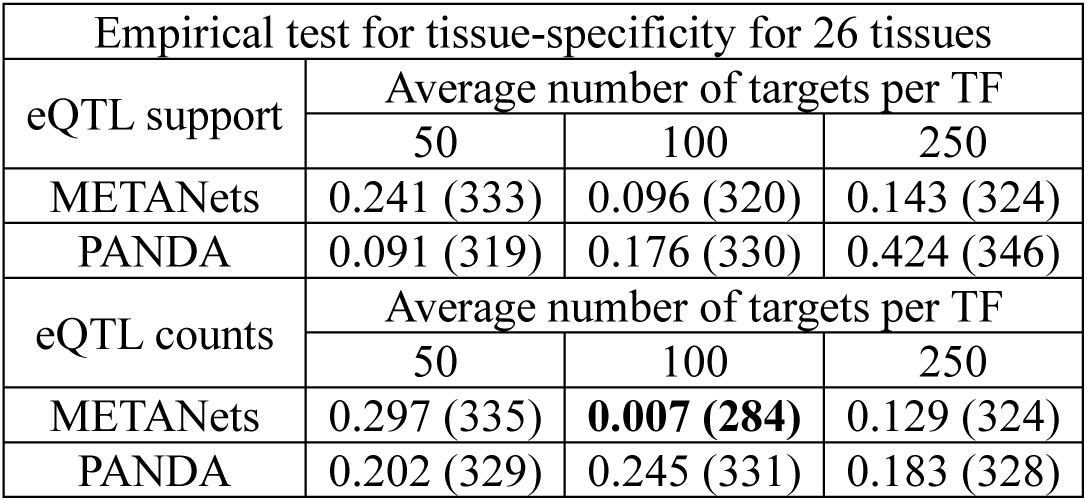
Tissue specificity evaluation of METANets and PANDA against the null across 26 tissues. Permutation test p-values (and the sum of observed matching ranks) against the distribution of 10000 permutation sums. METANets and PANDA networks did not show significant tissue specificity in eQTL support. METANets were significantly tissue specific in eQTL counts at the average 100 targets per TF threshold. PANDA networks remained not significantly tissue specific in eQTL counts. See also Supplementary Fig. S1.

| Empirical test for tissue-specificity for 26 tissues |  |  |  |
| --- | --- | --- | --- |
| eQTL support | Average number of targets per TF |  |  |
|  | 50 | 100 | 250 |
| METANets | 0.241 (333) | 0.096 (320) | 0.143 (324) |
| PANDA | 0.091 (319) | 0.176 (330) | 0.424 (346) |
| eQTL counts | Average number of targets per TF |  |  |
|  | 50 | 100 | 250 |
| METANets | 0.297 (335) | **0.007 (284)** | 0.129 (324) |
| PANDA | 0.202 (329) | 0.245 (331) | 0.183 (328) |

**Table 3.** Tissue specificity evaluation against Marbach. Wilcoxon signed-rank test p-values (and W statistics) between matching tissue ranks of METANets and Marbach networks for 36 tissues. Across all thresholds of average number of targets per TF, METANets and Marbach networks were not significantly different.

| Wilcoxon signed-rank test vs. Marbach |  |  |  |
| --- | --- | --- | --- |
| Average number of targets per TF | 50 | 100 | 250 |
| eQTL support | 0.95 (294.0) | 0.30 (266.5) | 0.88 (289.0) |
| eQTL counts | 0.24 (243.0) | 0.84 (269.5) | 0.43 (282.0) |

### 3.8 METANets Comparable to PANDA in Tissue Specificity but Outperform in Network Quality

Against the PANDA networks (26 tissues), neither PANDA nor METANets showed significant tissue specificity in eQTL support (Table 2). METANets were significantly better than chance in eQTL count in one threshold (average 100 targets per TF), but PANDA was not. The Wilcoxon signed-rank test found no significant difference between METANets and PANDA in either metric, indicating comparable tissue specificity (Table 4).

**Table 4.** Tissue specificity evaluation against PANDA. Wilcoxon signed-rank test p-values (and W statistics) between matching tissue ranks of METANets and PANDA networks for 26 tissues. Across all thresholds of average number of targets per TF, METANets and PANDA networks were not significantly different.

| Wilcoxon signed-rank test vs. PANDA |  |  |  |
| --- | --- | --- | --- |
| Average number of targets per TF | 50 | 100 | 250 |
| eQTL support | 0.57 (152.0) | 0.78 (152.0) | 0.87 (156.5) |
| eQTL counts | 0.90 (145.5) | 0.41 (132.0) | 0.88 (144.5) |

It is important to note that PANDA was originally developed as a general framework for inferring regulatory networks by integrating motif, expression, and protein-protein interaction data (Glass, et al., 2013). In its original form, PANDA produces a fully connected, weighted TF-gene network, but does not explicitly model tissue-specificity. The concept of *tissue-specific* PANDA networks was introduced by Sonawane et al. (2017), who applied PANDA to 38 GTEx tissues and then post-processed the resulting networks with a tissue specificity filtering step. In this approach, each edge’s weight in a given tissue was compared to its distribution across all tissues, and a normalized interquartile range (IQR) score was used to identify edges unusually strong in one tissue. Edges with high IQR scores were retained as “tissue specific.”

To enable direct comparison with the tissue-specific filtering strategy of Sonawane et al., we applied the same IQR-based approach to METANets. These filtered networks were evaluated without thresholding for top-scoring edges, as the number of TFs, genes, and edges varied drastically across tissues. Tissue-specific METANets showed significant eQTL support but not eQTL counts, while tissue-specific PANDA networks showed no significant signal in either metric (Table 5). Again, no significant difference was found between tissue-specific versions of METANets and PANDA networks (Wilcoxon signed-rank P > 0.05) (Table 6).

**Table 5.**
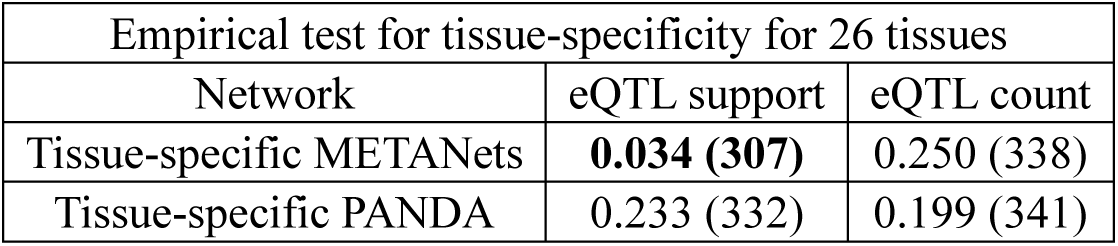
Empirical test of tissue-specific METANets and tissue-specific PANDA against the null across 26 tissues. Permutation test p-values (and the sum of observed matching ranks) against the distribution of 10000 permutation sums. Tissue-specific METANets showed significant tissue specificity in eQTL support and tissue-specific PANDA networks were not significantly tissue-specific. Neither tissue-specific METANets nor tissue-specific PANDA networks were significantly tissue-specific in eQTL counts. See also Supplementary Fig. S1.

| Empirical test for tissue-specificity for 26 tissues |  |  |
| --- | --- | --- |
| Network | eQTL support | eQTL count |
| Tissue-specific METANets | **0.034 (307)** | 0.250 (338) |
| Tissue-specific PANDA | 0.233 (332) | 0.199 (341) |

**Table 6.**
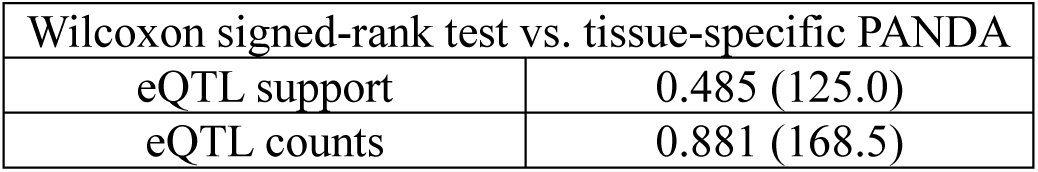
Tissue specificity evaluation against tissue-specific PANDA. Wilcoxon signed-rank test p-values (and W statistics) between matching tissue ranks of tissue-specific METANets and tissue-specific PANDA networks for 26 tissues. Full tissue-specific networks were evaluated, without thresholding for top-scoring edges. No significant difference between tissue-specific METANets and tissue-specific PANDA in both eQTL support and eQTL count.

| Wilcoxon signed-rank test vs. tissue-specific PANDA |  |
| --- | --- |
| eQTL support | 0.485 (125.0) |
| eQTL counts | 0.881 (168.5) |

Despite similar performance in tissue specificity, tissue-specific METANets outperformed tissue-specific PANDA in the Binding and GO-directness metrics (P<0.05 and P<0.001, respectively), while performing similarly in the GO metric (Fig. 4A-C). Tissue-specific METANets also performed better in the PPI metric overall (Fig. 4D).

**Fig. 4.**
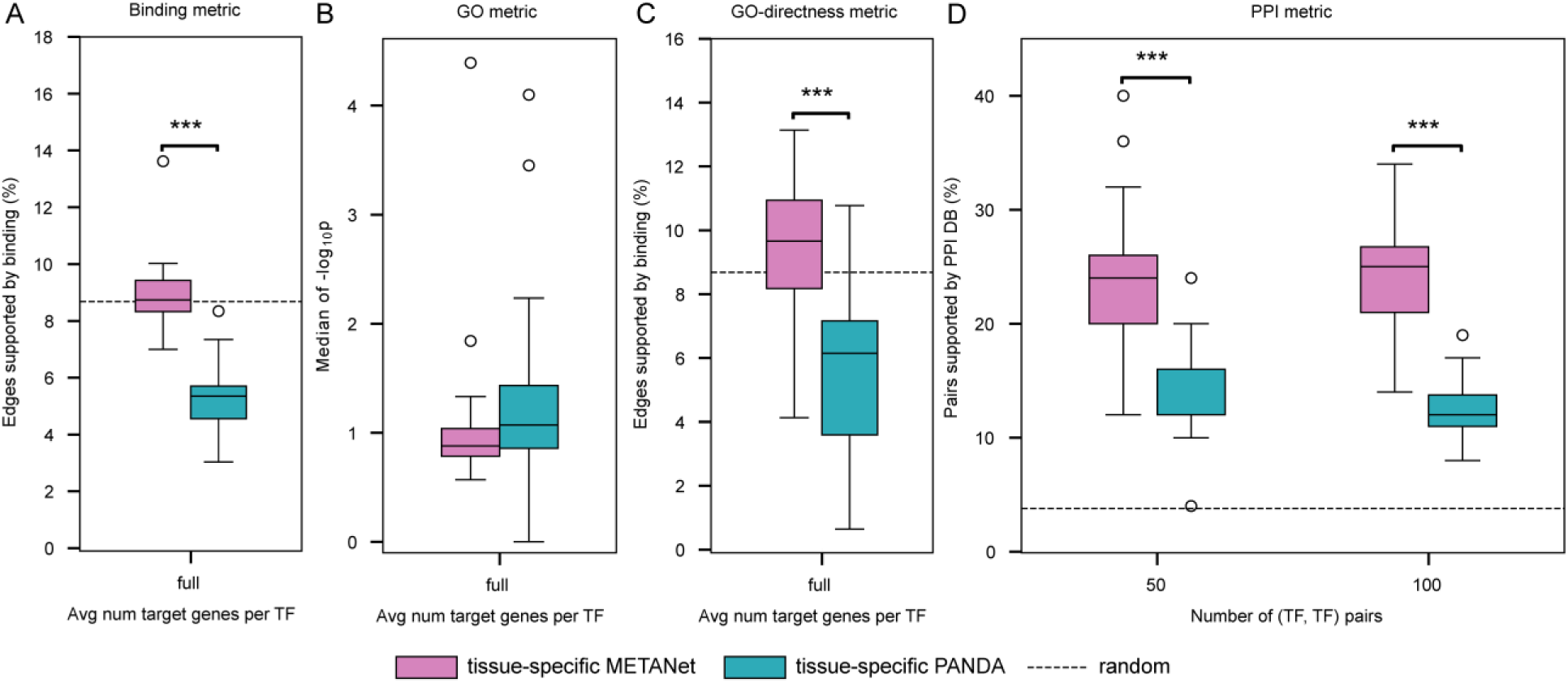
Network quality evaluation of tissue-specific networks. (**A-D**) Network quality evaluation of tissue-specific METANets and tissue-specific PANDA networks. (**A**) Binding metric. Tissue-specific METANets significantly outperformed tissue-specific PANDA networks (paired *t*-test, P<0.05). (**B**) GO metric. No significant difference between tissue-specific METANets and tissue-specific PANDA networks. (**C**) GO-directness metric. Tissue-specific METANets outperformed tissue-specific PANDA networks (P<0.001). (**D**) Tissue-specific METANets outperformed tissue-specific PANDA networks at thresholds of 50 and 100 (TF, TF) pairs. \**P≤*0.05, \*\**P≤*0.01, \*\*\**P≤*0.001.

### 3.9 Tissue-specific METANets are Useful for Genetic Analyses

To evaluate the biological relevance of tissue-specific METANets, we applied FISHNET, a method that uses prior biological knowledge to identify genes with suggestive association signals that are more likely to replicate across independent datasets (Acharya, et al., 2025). FISHNET integrates gene-level P-values with networks and functional annotations, based on the principle that spurious signals are randomly scattered, while true signals tend to cluster in functionally coherent subnetworks. Combining network topology, functional enrichment, and permutation-derived thresholds, FISHNET identifies a focused set of exceptional candidates termed FISHNET genes.

We used the whole blood tissue-specific METANet map as the reference network input to FISHNET. This network was partitioned into modules using a network modularization algorithm based on modularity optimization. As input, we provided gene-level P-values derived from measured TWAS of 11 traits associated with cardiovascular risk in the Long Life Family Study (LLFS) cohort (Acharya, et al., 2024) and performed replication analysis in the Framingham Heart Study (FHS) cohort (KANNEL, et al., 1979; Splansky, et al., 2007). Across the 11 traits, FISHNET identified 114 FISHNET genes of which 24 were replicated. Using the whole blood tissue-specific PANDA network as input, we identified 0 FISHNET genes (see Supplementary Tables S1 and S2).

A central biological story emerging from these results centers on *SREBF2*, a replicated FISHNET gene negatively associated with HDL. Our findings support a model in which high HDL levels may suppress *SREBF2* activity, whereas low HDL permits *SREBF2* activation and sustains inflammatory signaling. *SREBF2* encodes SREBP2, a master transcription factor of sterol metabolism, canonically activated in adipocytes under low-cholesterol conditions (Bauer, et al., 2011; Horton, et al., 2002; Jiang, et al., 2005). Recent studies have shown that inflammatory stress can also activate SREBP2, enabling it to bind and induce inflammatory and interferon-response genes (Fowler, et al., 2023; Kusnadi, et al., 2019; Zhao, et al., 2011). In the whole blood tissue-specific METANet map, *SREBF2* appears in the same module as other negatively HDL-associated FISHNET genes–*S100A9, S100A12, TLR8,* and *ITGB2*–all of which converge on pathways that activate the NADPH oxidase 2 (NOX2) complex. *S100A9* and *S100A12* are calgranulins, a type of alarmin, that can act as ligands for the receptor for advanced glycation end products (RAGE), triggering assembly and activation of the NOX2 complex (Bowman and Schmidt, 2011; Dong, et al., 2022; Xiao, et al., 2020). *TLR8* is a pattern recognition receptor that primes NOX2 activation in human neutrophils (Makni-Maalej, et al., 2015). *ITGB2* promotes leukocyte adhesion to the endothelium and triggers “outside-in” signaling that further promotes NOX2 assembly and activation (Chen, et al., 2016). Collectively, these genes link *SREBF2* to NOX2-based excessive production of reactive oxygen species (ROS), which oxidize HDL and convert it into a pro-inflammatory form (oxHDL) that contributes to atherosclerosis. Notably, *S100A9*, *S100A12*, and *TLR8* also activate NF-κB, inducing cytokines (TNFα and IL-1β) known to activate *SREBF2* (Cluzeau, et al., 2017; Qin, et al., 2006; Yang, et al., 2007; Zhang, et al., 2023). Together, these findings suggest a positive feedback loop in which reduced HDL allows *SREBF2* activation, which in turn promotes inflammatory cascades that further lower HDL, providing a mechanistic link between cholesterol metabolism, immune activation, and chronic low-grade inflammation characteristic of obesity.

## 4 DISCUSSION

In this work, we developed the METANet framework, a supervised ensemble approach that combines TF motifs, cis-regulatory element (CRE) activity, and expression-derived features to construct tissue-specific TF network maps across 36 human tissues. Our modeling approach relied on three fundamental ideas: (1) using gene expression data (functional) to predict TF binding events (direct) prioritizes direct, functional targets of TFs, (2) incorporating both tissue-aggregate and tissue-specific expression features captures both the shared and tissue-specific regulatory edges, and (3) using tree-based regression models to incorporate both linear and nonlinear expression patterns improves model robustness. Our evaluations across multiple orthogonal metrics show that our approach significantly outperforms motif-only, expression-only, and unsupervised integrative methods in capturing direct and functionally coherent TF-TG relationships.

Gene expression-based models are adept at identifying functionally coherent gene sets, but the expression of a TF and a potential target gene could be correlated for reasons other than the TF regulating the gene directly: the TF might regulate indirectly (mediated by other proteins), or the TF and the gene might simply be co-regulated by a third factor. Conversely, motif-based approaches anchor predictions in physical binding potential but cannot confirm functional relevance, as potential binding sites are frequently unoccupied (Dror, et al., 2015; Wang, et al., 2012) and even occupied sites often do not affect transcription (Cusanovich, et al., 2014; Kang, et al., 2020; Mahendrawada, et al., 2025; Slattery, et al., 2014; White, et al., 2013). By training our models to explicitly predict TF ChIP-seq binding events, METANet leverages the strengths of both data types without a noticeable tradeoff. Our results demonstrate the resulting METANet maps (METANets) consistently outperforming the motif-only Marbach networks and our expression-only ETANets in the GO-directness metric that jointly assess directness and functional coherence.

The METANet framework also illustrates the power of incorporating diverse feature types to model the complexity of gene regulation. We found that regulatory relationships are not exclusively linear or non-linear; by combining features from both LASSO (linear) and BART (non-linear) regression models, METANets captured a more direct and functionally coherent set of TF targets. Similarly, we found that neither tissue-specific nor tissue-aggregate features clearly outperformed the other in isolation, but together they predicted more robust sets of TF targets. The METANet approach performed best in the individual Binding and GO metrics without sacrificing GO-directness as a tradeoff. The practical utility of this approach was highlighted in our downstream genetic analysis, where the whole blood METANet map enabled FISHNET to identify replicated gene-trait associations, proposing a module linking *SREBF2* to HDL and chronic inflammation.

Our work presents a subtle methodological shift for the field of regulatory network inference. For years, the community has largely relied on two distinct categories of methods: those based on physical binding evidence like TF motifs, which often lack functional validation, and those based on gene expression, which are susceptible to confounding by indirect relationships. While unsupervised integration methods have attempted to bridge this gap, the METANet approach presents a new alternative by successfully implementing a supervised learning framework. The core innovation is reframing the problem: instead of merely correlating different data types, we use functional data (gene expression) to predict direct physical interactions (TF binding).

This approach provides a methodological foundation for using the ever-expanding space of omics data integration. The principle of using one data modality as a ground-truth label to train predictive models on other, more widely available data types can be generalized beyond the specific application in this work. It makes TF network mapping more broadly applicable, as ChIP-seq data for every TF is not available, while RNA-seq data is abundant. Our approach can make use of ChIP-seq in predicting targets of TFs that have ChIP-Seq data, but it can also extract general patterns in gene expression data that are predictive of direct binding and use those patterns to infer targets for TFs that do not have ChIP-Seq data. We also presented a new evaluation of networks based on utility for applications rather than network characteristics.

Several limitations suggest avenues for further development. One key limitation is the lack of tissue-specific TF binding data in our framework. While tissue-specific expression data are readily available at scale, obtaining comparable TF binding data across tissues remains challenging. As a result, METANet relied on tissue-aggregated TF binding data. Emerging paired-seq technologies that jointly measure TF binding and gene expression in the same cell could make predictions of TF binding events more robust and the single-cell measurements can facilitate the direct mapping of granular cell-type-specific networks. Development of scalable methods to address the inherent sparsity of single-cell data will be crucial for realizing this goal. Another opportunity lies in extending the METANet framework to additional omics data types. Chromatin accessibility (ATAC-seq) and chromatin conformation (Hi-C, HiChIP) represent rich sources of contextual information that could further refine predictions of TF-TG relationships. Beyond tissues, the same framework could be adapted to disease states, environmental exposures, or dynamic perturbations, producing condition-specific network maps that illuminate regulatory mechanisms underlying phenotypic variation.

In summary, METANet advances the state of regulatory network mapping by demonstrating that supervised integrative approaches can accurately prioritize direct, functional TF-TG interactions. Beyond their immediate utility, METANet highlights the potential of combining heterogeneous data sources to produce context-dependent regulatory maps, setting the stage for increasingly precise models of human gene regulation.

## DATA AND CODE AVAILABILITY

All data and code are available at https://doi.org/10.5281/zenodo.17309371.

## ACKNOWLEDGEMENTS

We are grateful to the entire Long Life consortium, its participants, and its investigators, without whom this work would not have been possible. The Framingham Heart Study is conducted and supported by the National Heart, Lung, and Blood Institute (NHLBI) in collaboration with Boston University [N01-HC-25195, HHSN268201500001I, 75N92019D00031]. This manuscript was not prepared in collaboration with investigators of the Framingham Heart Study and does not necessarily reflect the opinions or views of the Framingham Heart Study, Boston University, or NHLBI.

