## Supplementary Material for "METANet: A supervised ensemble learning framework for reconstructing direct and functional tissue-specific transcription factor networks"

Michael R. Brent

(314) 362-7238

Campus Box 8510

Washington University

Saint Louis, MO 63130

### Supplementary Figure S1

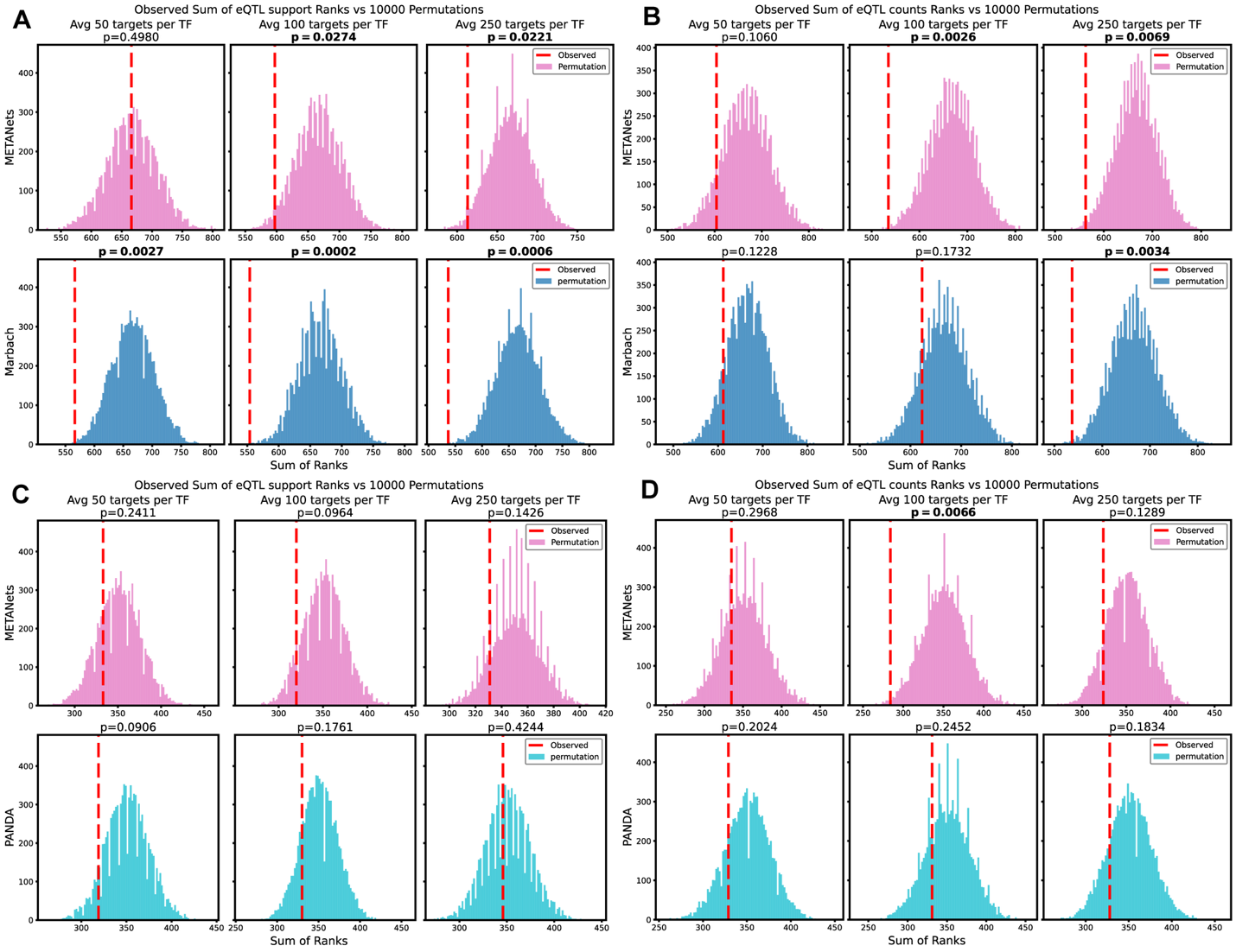

**Supplementary Fig S1 Tissue specificity evaluation against the empirical null.** The sum of eQTL support or eQTL counts ranks (dashed red line) plotted against a null distribution of 10000 sums of permuted ranks. METANets and Marbach networks were evaluated across 36 tissues for (**A**) eQTL support and (**B**) eQTL counts. METANets and PANDA networks in evaluated across 26 tissues for (**C**) eQTL support and (**D**) eQTL counts. METANets have tissue specificity comparable to other networks, except that Marbach networks have better eQTL support at average 50 targets per TF (panel A).

### Supplementary Figure S2

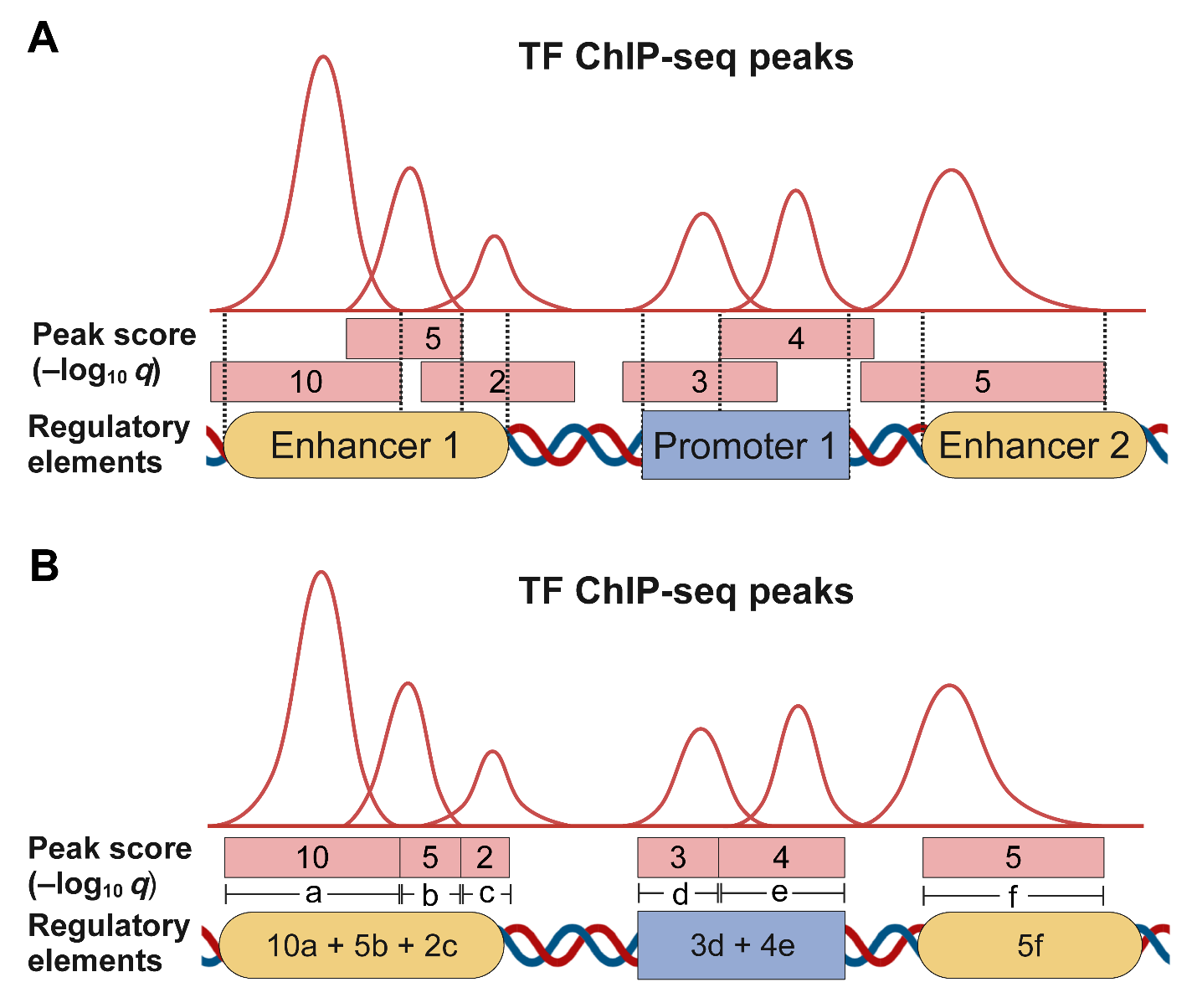
 **Supplementary Fig S2 Example illustration of assigning TF binding scores to FANTOM5 regulatory elements.** (A) Multiple TF ChIP-seq peaks may overlap in a single regulatory element. Dotted vertical lines indicate cutoffs for overlapping peaks, retaining the peak with maximum peak. (B) Each regulatory element’s score is calculated as the sum of retained peak scores weighted by the peak region length.

### Supplementary Figure S3

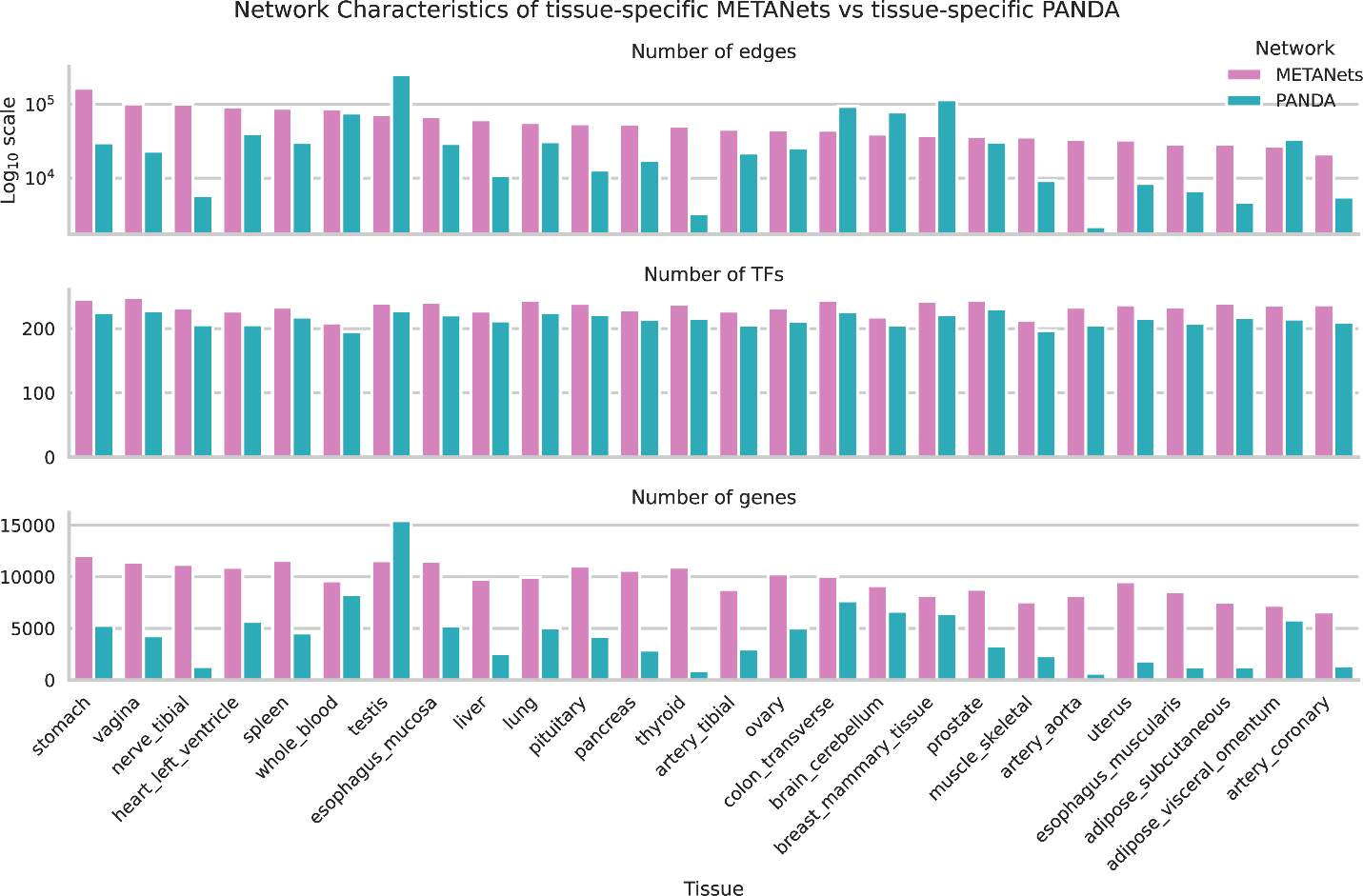

**Supplementary Fig S3 Network characteristics of tissue-specific METANets and tissue-specific PANDA networks** PANDA networks were first restricted to protein-coding genes, with self-regulatory edges excluded, and subsequently subsetted to TFs present in the corresponding tissue’s METANet. See Additional file 2: Table S4 for tabulated details of this figure.

### Supplementary Figure S4

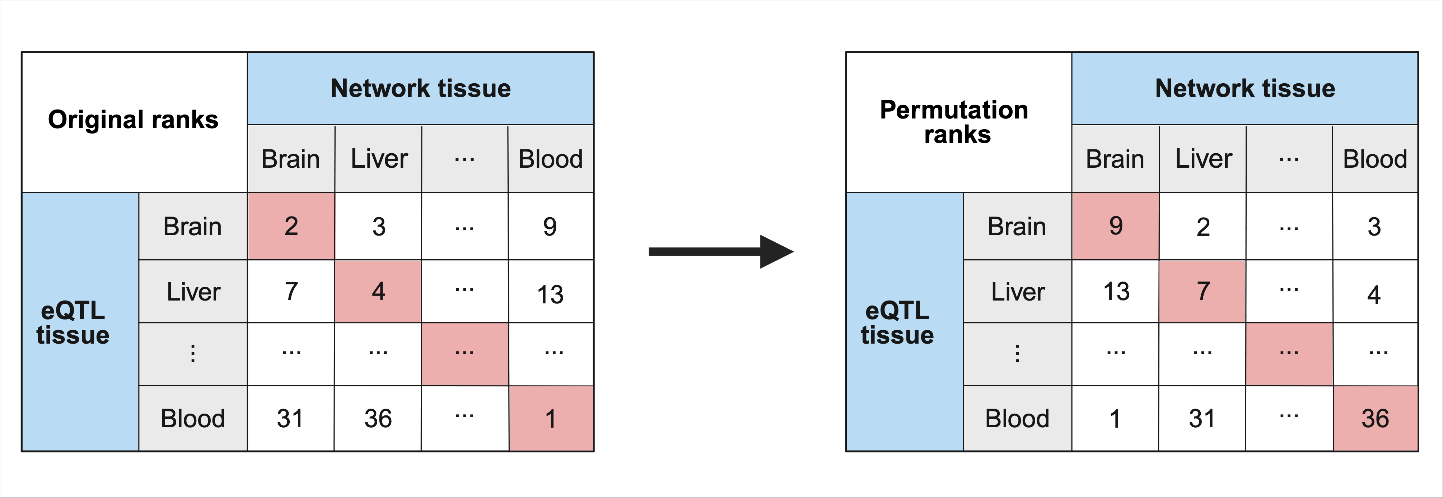

**Supplementary Fig S4 Permutation to generate empirical null distribution of ranks** For each tissue’s eQTL data, all tissue networks are ranked based on either the eQTL support or eQTL counts metric, breaking ties using the other metric. Each diagonal entry in the original ranks table represents the rank of the “matching” network for the given tissue’s eQTL. The sum of the diagonal entries in the original ranks table (left) is the observed statistic. For each of 10000 permutations, we shuffle the columns of the table. The sum of the diagonal entries in this shuffled table is the permuted statistic. This process simulates the null hypothesis that there is no true correspondence between a network and the tissue it was constructed for.

### Supplementary Methods

#### Mapping binding peaks to regulatory regions

For each TF, all high-confidence (FDR q-value $\leq$ 0.01) binding peaks within FANTOM5 promoter and enhancer regions were examined (Andersson, et al., 2014; The GTEx Consortium, 2020). In cases where multiple overlapping peaks were present within a single regulatory element, the peak with the highest confidence score among the overlapping regions was retained. A score was calculated for each regulatory element by summing the confidence scores of the retained peaks, weighted by their length in base pairs (see Supplementary Fig. S2). For each gene, a gene score was derived by aggregating the scores of all regulatory elements annotated for the gene. Since TF binding events within each regulatory region represent independent evidence of gene regulation, the aggregated score across regulatory elements reflects the cumulative likelihood of the TF regulating the gene.

#### Construction Of Binding Labels

The network mapping approach uses binary labels representing binding events for each TF-gene pair. We assigned a positive label to genes with nonzero binding scores for each TF to construct binary labels for TF-gene binding events. If more than 10% of genes had a positive score for a given TF, only the top 10% of highest-scoring genes were retained as positive labels. All remaining gene pairs were labeled as negative.

#### Processing of PANDA Networks

For benchmarking, we downloaded PANDA networks for 38 human tissues from Zenodo <https://doi.org/10.5281/zenodo.838734> (Sonawane, 2017). To enable direct comparison with our METANets, we retained 26 tissues shared with the 36 METANet tissues. For each tissue, we used two versions of the PANDA networks: (1) the full networks containing weighted edges between all transcription factors (TFs) and genes, and (2) the tissue-specific networks provided in the same dataset, in which tissue-specific edges had already been annotated by the original authors. To maintain consistency, both versions of the PANDA networks were subjected to the same downstream processing steps: restricting to protein-coding gene targets, removing self-regulatory edges, and retaining only the subset of TFs common to both PANDA and METANets.

#### Tissue-specific network filtering

Tissue-specific regulatory edges were identified using the filtering approach originally introduced by Sonawane et al. (Sonawane, et al., 2017). For each TF-TG pair *(i,j)*, the edge weight in tissue *t* is denoted $w_{ij}^{(t)}$. For each edge, we computed a normalized specificity score by comparing the tissue-specific weight to the distribution of weights across all tissues:

$$s_{ij}^{\left( t \right)}=\frac{w_{ij}^{(t)}-\text{median}(w_{ij})}{\text{IQR}(w_{ij})},$$

where $\text{median}(w_{ij})$ and $\text{IQR}(w_{ij})$ are calculated across all tissues for the same edge. Edges with $s_{ij}^{\left( t \right)}>2$ were considered specific to tissue *t*. This filtering approach was applied to METANets to construct tissue-specific versions of our networks, directly comparable to the tissue-specific PANDA networks generated in Sonawane et al.

Each METANet map varies in total edge count due to differences in gene expression coverage; some genes fail to pass tissue-specific expression filters and thus lack corresponding edges in certain tissues. Because the tissue-specificity score $s_{ij}^{(t)}$ depends on computing both the median and the IQR of edge weights $w_{ij}$ across all tissues, edges missing in many tissues results in too few observations to reliably estimate their across-tissue variability. We therefore excluded edges absent in more than 12 of the 36 tissues. This step retained 77.16% of the complete edge set.

#### Validation of Tissue Specificity using eQTL Data

We downloaded cis-eQTL data from the GTEx consortium V8 release (The GTEx Consortium, 2020). Significant variant-gene associations were obtained for each of the 36 tissues. For each TF-target gene (TF-TG) edge in a network, we determined whether one or more of a tissue’s significant variants overlapped the TF’s ChIP-seq peaks within the promoter or annotated enhancers of the target gene. We defined two complementary metrics for a given network:

1. eQTL support: the fraction of edges supported by at least one tissue-specific eQTL.
2. eQTL count: the sum, across all edges, of the number of tissue-specific eQTLs supporting each edge.

To assess a network’s tissue-specificity, we performed a permutation test. First, we calculated an observed statistic $S_{obs}$ by ranking all 36 networks within each tissue and summing the ranks of the “matching” pairs (e.g., the rank of the liver network in the liver tissue). A smaller sum indicates better overall specificity. Next, to generate a null distribution, we performed 10000 permutations. In each permutation, we randomly shuffled the network labels before summing the ranks of the newly assigned matching pairs to obtain the permutation statistic $S_{perm}$. This process simulates the null hypothesis that there is no true correspondence between a network and its native tissue. The empirical p-value was then calculated as the proportion of permutations with $S_{perm}\leq S_{obs}$ (see Supplementary Fig. S4).

Direct comparisons between different networks were performed using the Wilcoxon signed-rank test on the distributions of matching ranks.

#### Module Identification for FISHNET

We applied FISHNET, a network-guided replication framework that integrates prior biological knowledge with gene-level association statistics, to evaluate the biological relevance of tissue-specific METANets (Acharya, et al., 2025). FISHNET is designed to prioritize genes that exhibit suggestive association signals but are more likely than expected by chance to replicate across independent datasets.

The primary input to FISHNET was the whole blood tissue-specific METANet. For comparison, we also applied FISHNET using the whole blood tissue-specific PANDA network constructed by Sonawane et al. (2017). Network modules were identified using the top three algorithms from the Disease Module Identification, DREAM Challenge (Choobdar, et al., 2019): K1 (kernel clustering), M1 (modularity optimization), and R1 (random walk-based clustering). We implemented these algorithms through the MONET package (Tomasoni, et al., 2020) and applied each to the whole-blood-specific TF network maps from METANet and PANDA. The M1 algorithm was run in both directed and undirected modes by varying the “--linksdir” parameter.

As input gene-level signals, we used transcriptome-wide association study (TWAS) p-values derived from the Long Life Family Study (LLFS). We considered 11 traits associated with cardiovascular risk, including body mass index (BMI), high-density lipoprotein (HDL), low-density lipoprotein (LDL), and triglycerides, among others. Replication was assessed using gene-level association statistics from the Framingham Heart Study (FHS) cohort, analyzed for the same 11 traits. Details on RNA-seq data processing and covariate adjustments have been previously described by Acharya et al. (Acharya, et al., 2025).

Summary characteristics of the modules generated by each algorithm and network are provided in Additional file 2: Table S1. The final modules used for the *SREBF2*-related biological analysis were derived from the directed M1 algorithm.
